# Stem cell regulators control a G1 duration gradient in the plant root meristem

**DOI:** 10.1101/2022.03.09.483577

**Authors:** Clara Echevarria, Bénédicte Desvoyes, Marco Marconi, José Manuel Franco-Zorrila, Laura Lee, Masaaki Umeda, Robert Sablowski, Kenneth D. Birnbaum, Krzysztof Wabnik, Crisanto Gutierrez

## Abstract

In meristems, where new plant organs initiate, key stem cell regulators have been identified, but their link to cell cycle progression remains unclear. Here, we show that the root meristem has a positional gradient of G1 duration that ranges from ∼2 h near the meristem boundary to more than 20 h in stem cells and early derivatives. Mutants in the *PLETHORA* (*PLT*) genes shortened G1 length and flattened its gradient. Computer modeling of an incoherent feed-forward loop (IFFL) predicted the inference of a negative regulatory pathway. We propose that *PLT* genes play opposing roles, maintaining meristem and stem cell activity and inhibiting G1 progression through the CDK inhibitor KRP5, a PLT target, and RBR1. This establishes a previously undescribed proximal-distal feature of the root meristem in which a G1 duration gradient is shaped by stem cell and meristem maintenance regulators.

---

The production of new cells is required during organogenesis at the same time that patterning genes establish different organ domains and cell types. Thus, a fundamental challenge in cellular and developmental biology is to understand the coordination between cell patterning and cell division during organogenesis. In animals, the cell cycle phase in which inductive cues are received can dictate the choice of cell fate and the switch to pluripotency (Pauklin & Vallier, 2013; Boward *et al*, 2016). In plants, cell patterning decisions are integrated with cell division (Costa & Shaw, 2006; Meyer *et al*, 2017; Desvoyes & Gutierrez, 2020; Sablowski, R. & Gutierrez, C., 2021). However, the pathways linking cell cycle progression to spatio-temporal dynamics of upstream developmental cues are not well understood. In fact, the molecular basis of cell cycle phase progression in a developmental context is largely unknown. This lack of knowledge stems from the difficulties in measuring cell cycle phase parameters, in particular G1, which is a potential control point for cell fate, at single-cell resolution in a developing organ.

To address this question, we focused on the root apical meristem (RAM) of *Arabidopsis thaliana* (Fig. S1), since it is amenable for live-imaging and it possesses a stereotyped anatomy, allowing us to visually track cell type files from stem cell to differentiated state (Dolan *et al*, 1993; Scheres, 2007). The PlaCCI Arabidopsis line (Desvoyes *et al*, 2020) is an ideal tool to measure directly the G1 duration since the CDT1a-CFP marker starts to accumulate soon after mitosis and is rapidly degraded at the G1/S transition.

The analysis of time-lapse images (Fig. 1A, B and Video S1) revealed that G1 duration was inversely correlated with the distance from the stem cell niche. G1 duration followed a gradient in all tissues analyzed (trichoblasts, atrichoblasts, cortex, endodermis) that smoothly changed from ∼2-4 h in cells towards the RAM boundary up to ≥20 h in the more distal half of the RAM, close to the stem cell niche (Fig. 1C).

**Fig. 1.**
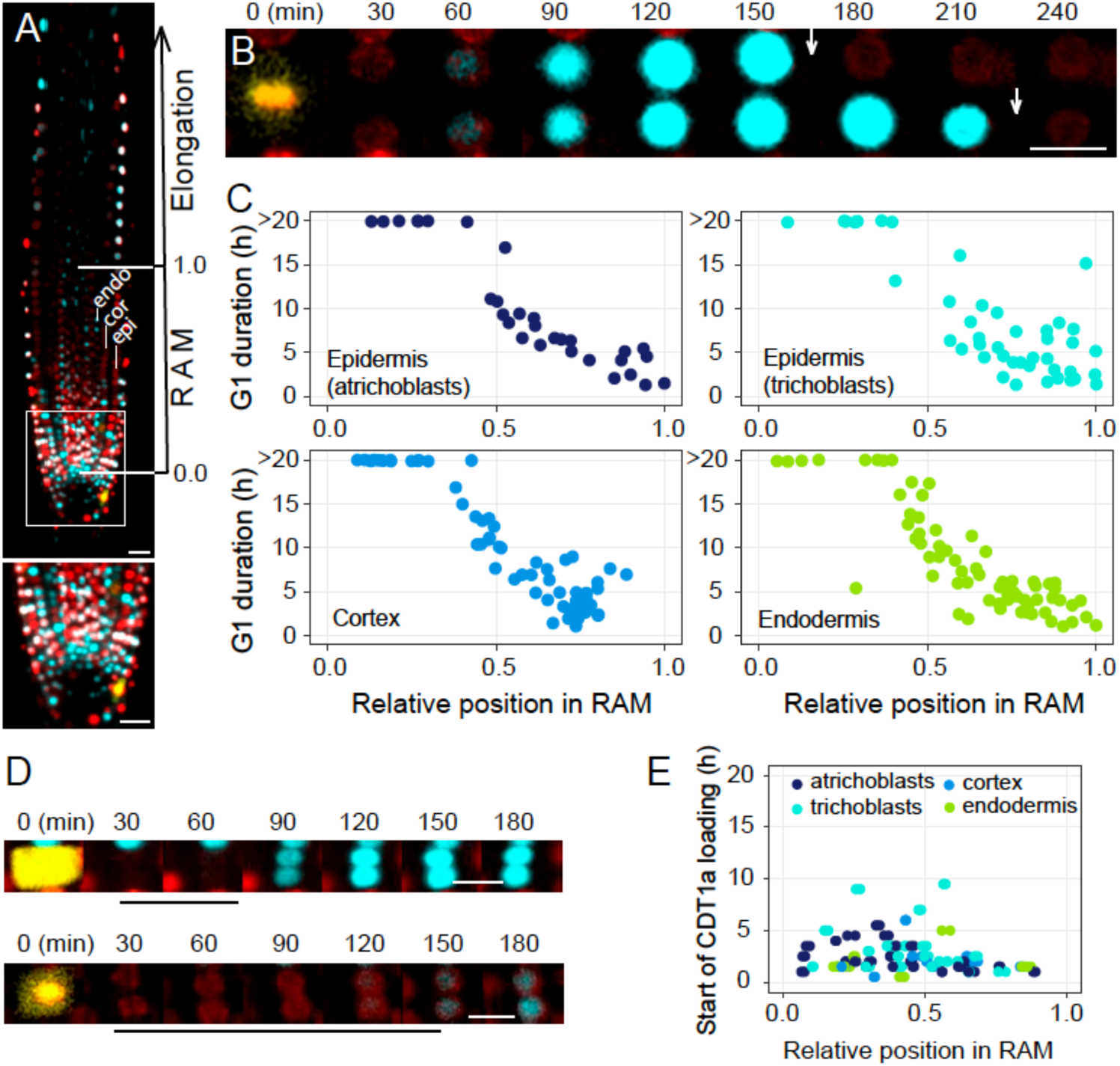
Root stem cells and early derivatives develop very long G1. **A**, In the upper panel, a root tip of the three-color cell cycle marker line PlaCCI showing nuclei in G1 (cyan), S + early G2 (red) and late G2 + M (yellow). The image includes the root apical meristem (RAM) at the tip, rich in cells, followed by the elongation zone in the upper part of the root. The inset (white square) is enlarged in the bottom panel focusing on the quiescent center (QC), the stem cell niche and the surrounding early derivatives. The epidermal (epi), which contains atrichoblasts and trichoblasts, cortical (cor) and endodermal (endo) layers are indicated. Note that the epidermis contains both atrichoblasts and trichoblasts). The position of the quiescent center (QC) is set as 0.0 and the RAM boundary as 1.0. Bar = 20 µm. **B**, Live-imaging showing a trichoblast, located at position 0.76 of the RAM, in metaphase (0 min). After division, the two daughter cells load CDT1a (cyan). Since CDT1a is rapidly degraded at the G1/S transition (vertical arrows), a process showing some variability between daughter cells, it is a proxy of the G1 duration. Scale bar = 10 µm. **C**, G1 duration in four root cell types (atrichoblasts, trichoblasts, cortex, endodermis), as indicated, along the root apical meristem (RAM). The position of the QC is set as 0.0 and the RAM boundary as 1.0. Data points correspond to the G1 duration of individual cells quantified from different recorded videos (up to 20h long using confocal microscopy). Data points correspond to the G1 duration of individual cells quantified from different recorded videos (up to 20h long using confocal microscopy). The number of cells recorded were 37, 46, 68, and 76 for atrichoblasts, trichoblasts, cortex and endodermis, respectively. **D**, Live-imaging of cells showing the period from anaphase until initiation of CDT1a loading. Two examples of a short (upper panel) and a long (lower panel) period of CDT1a loading initiation (black line) are shown. Scale bar = 10 µm. **E**, Quantification of CDT1a loading time (between anaphase and the first detectable CDT1a signals) along the RAM in four cell types, as indicated. Total n = 128 cells.

It is generally assumed that cell cycle duration is constant in the transit amplifying compartment that constitutes most of the RAM (Ivanov & Dubrovsky, 1997; Fiorani & Beemster, 2006; Pacheco-Escobedo *et al*, 2016). Some recent reports suggest a gradient of cell cycle duration from the stem cells towards the RAM boundary (Rahni *et al*, 2016; Rahni & Birnbaum, 2019). Our results showed that the G1 duration was generally longer in most cells in the distal RAM than the average cell cycle values reported, considering a constant cell cycle along the RAM (Zhukovskaya *et al*, 2018).

In one of the few attempts to measure cell cycle phase duration along the meristem, early studies with irradiated root cells found a long G1 duration in cells around the QC (Clowes, 1965). However, using the PlaCCI marker line we found that G1 duration is organized in a proximal-distal gradient (Fig. 1C). Thus, not only the stem cells but also their early derivatives (up to ∼1/3 of the RAM) develop considerably much longer G1 phases than rapidly cycling cells in the more shootward half of the RAM. Furthermore, since the longitudinal axis of RAM is correlated with the trajectory of root cells, our results reveal that the G1 duration is developmentally regulated along the RAM, showing a distinct G1 gradient along the proximal-distal root axis.

To check whether our G1 duration measurements were not biased by the appearance of the CDT1a signal, we determined the accumulation kinetics of CDT1a in early G1 (time between anaphase and the first detectable CDT1a signal) along the RAM (Fig. 1D). This analysis revealed that the differences in all cell types analyzed were relatively small compared with the total G1 length, insufficient to explain the G1 duration gradient along the longitudinal RAM axis (Fig. 1E). Also, although variability exists in CDT1a accumulation, it does not correlate with a proximal-distal gradient. Therefore, we conclude that G1 phase duration of stem cells and their derivatives within the proliferation domain of the RAM is dictated by a slow G1 progression and not merely a slower kinetics of CDT1a accumulation.

We also sought to determine if other phases of the cell cycle offset the G1 gradient to keep the entire cell cycle constant, at least in the transit amplifying zone of the root. One possibility is that an increasing gradient of delayed or prolonged G2 progression along the meristem offsets the increasingly rapid G1 duration. We measured the average G2 duration by quantifying the appearance of labeled mitosis in an EdU pulse-chase experiment (Fig. S2) (Echevarría *et al*, 2021) and found significant, though small, differences between the external (∼4.5h) and internal (∼3h) cell layers (Fig. S3). We also measured G2 duration of individual cells along the RAM by recording positional information of EdU-labeled mitosis (Fig. 2A). We noticed that EdU-labeled mitosis tended to appear earlier in the distal half of the RAM (∼40% the total distance between the QC and the RAM boundary), indicating a slightly faster G2 in the distal half of the RAM compared to the proximal half (Fig. 2B), consistent with previous results obtained for the epidermis (Otero *et al*, 2016). Time-lapse live imaging to measure the kinetics of CYCB1;1 accumulation and degradation in individual cells along the RAM revealed that the extent of detectable CYCB1;1 showed a certain degree of variability (Fig. 2C, D). However, the variation was insufficient to compensate for the G1 differences. Thus, G2 duration does not offset and compensate the large variation in G1.

**Fig. 2.**
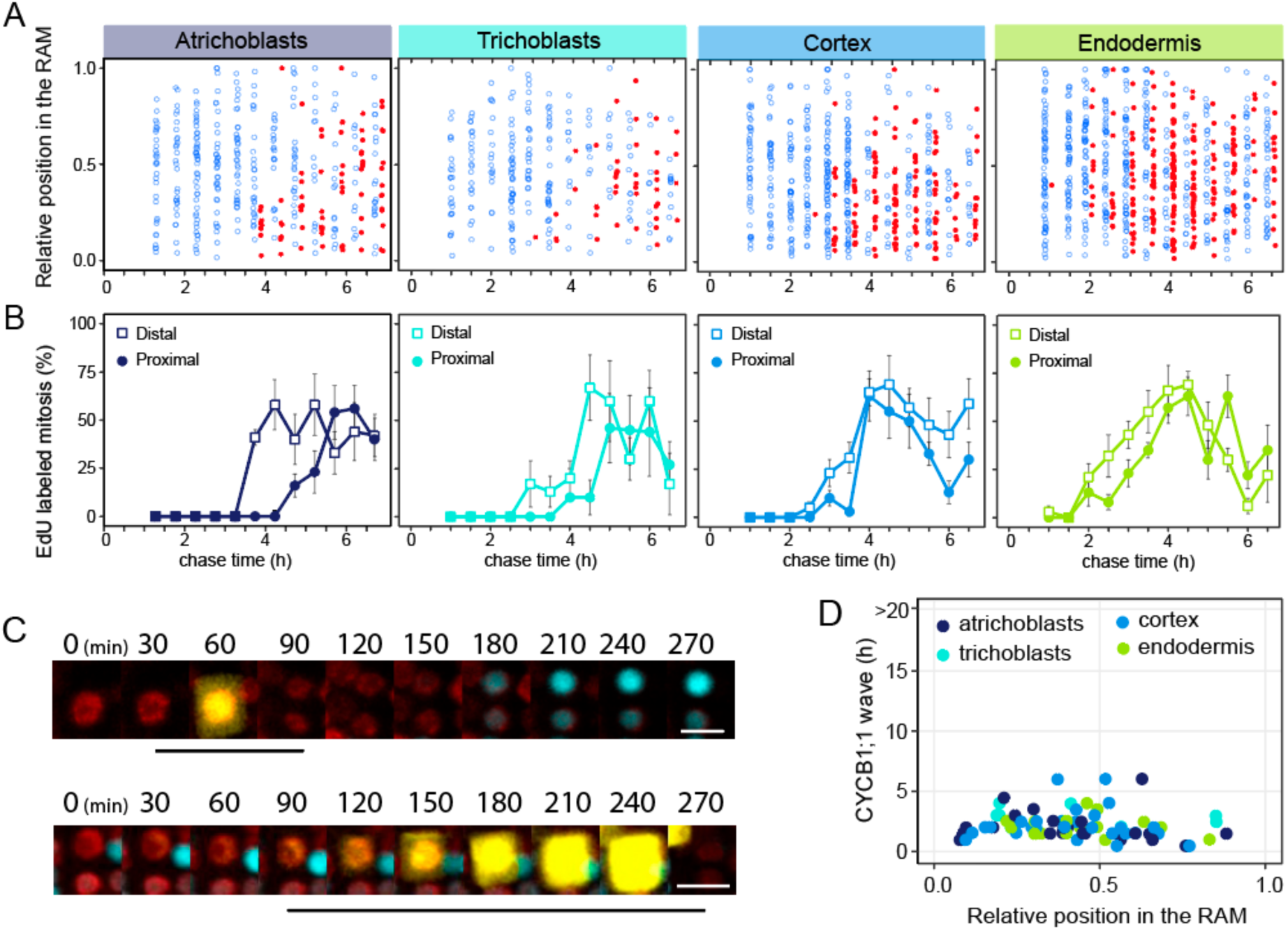
Differences in G2 duration do not compensate for the long G1. **A**, Analysis of G2 progression by measuring the appearance of mitotic cells labeled after an EdU pulse (20 min). Unlabeled (blue dots) and labeled (red dots) mitosis were scored at various chase times after the EdU pulse, according to their position along the RAM (0 = QC, 1.0 = RAM boundary, as described in Fig. 1C). Quantifications were carried out separately for various cell types, as indicated. **B**, Quantification of EdU-labeled mitosis at various chase times in two regions of the RAM: distal (empty squares) corresponds to the region located between the position 0 (QC) up to 0.4 of the RAM; proximal (filled circles) includes the rest of the RAM (from 0.4 to its boundary at 1.0). Data are mean values ± s.d. The experiment was carried out in duplicate, processing 5 root meristems per biological replicate for each chase time. Total number of cells analyzed for each cell type were: atrichoblasts (n=312 EdU–, n=75 EdU+), trichoblasts (n=237 EdU–, n=42 EdU+), cortex (n=401 EdU–, n=139 EdU+), endodermis (n=495 EdU–, n=237 EdU+). **C**, Live-imaging of cells showing the dynamics of CYCB1;1 loading in late G2 and its degradation at the metaphase/anaphase transition. Two examples of cells with a short (upper panel; atrichoblast located at relative position 0.41 of the RAM) and long (lower panel; trichoblast at position 0.37) G2+prophase+metaphase period (black bars under each panel) are shown. Scale bar = 10 µm. **D**, Duration of the CYCB1;1 wave (time between first detectable signal and complete degradation of CYCB1;1 at the metaphase/anaphase transition) along the root meristem in four root cell types, as indicated. Total n = 75 cells.

We next aimed to identify the regulatory network underlying the G1 duration gradient. To this end, we analyzed transcriptomic datasets of root cell types (Brady *et al*, 2007) searching for genes displaying a variable expression pattern along the RAM that could show a relationship with the G1 gradient. An unbiased clustering of the 1,472 gene subset with a variable expression, using a weighted correlation network analysis (WGCNA) led us to identify highly correlated gene expression patterns (Table S1). The seven different patterns (M1 through M7) could be classified into 3 groups, corresponding to genes with high expression only in the RAM (type A), in the middle of the root (type B) or up in the differentiated zone of the root (type C; Fig. S4 and Table S1).

To identify putative transcriptional regulators that could contribute to the different expression profiles, we searched for enriched transcription factor binding sites (TFBS) in the promoters of genes in each module (Fig. 3A). Focusing on the M1 module, which has a profile similar to the G1 duration gradient, we identified a high score enrichment for binding sites of APETALA2-AINTEGUMENTA (AP2-ANT), bHLH and SQUAMOSA PROTEIN BINDING (SPB) TF family members.

**Fig. 3.**
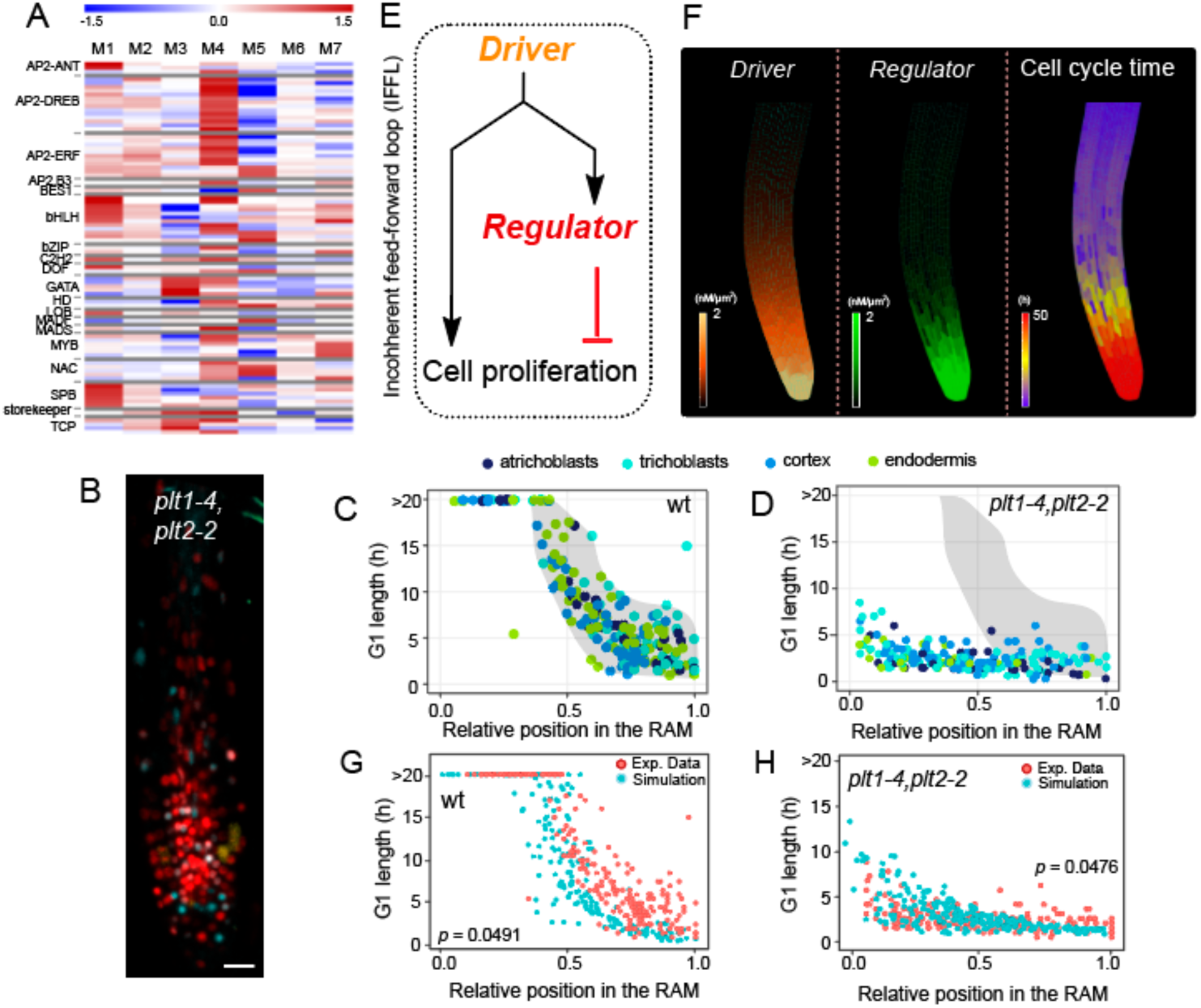
Spatio-temporal computer model of root cell proliferation for developmental control of G1 in wild type and loss of PLT function. **A**, Heat map showing enrichment of transcription factor binding sites (TFBS) in rows, in the gene sets from each module (M1-M7, columns). TFBS were mapped in 1 kb regions upstream of the transcriptional start sites with FIMO, and fold-enrichment in log2 scale calculated by mapping the same sites in the complete Arabidopsis promoter set. **B**, Root tip of a *plt1-4,plt2-2* double mutant showing the reduced size of the RAM. **C**, Summary of G1 duration values in the wild type, taken from Fig. 1C and included here to facilitate comparison with the *plt1-4,plt2-2* double mutant. The shaded shape includes most of the G1 values measured in the wild type. **D**, G1 duration in four root cell types (atrichoblasts, trichoblasts, cortex, endodermis; n = 36, 26, 81 and 64, respectively), as indicated by the color code, along the RAM in the *plt1-4,plt2-2* double mutant. The shaded shape is taken from Fig. 3C and represents the values obtained for the wild type. **E**, Schematic diagram of the incoherent feed-forward loop (IFFL) mechanism underlying the model. The diagram shows the main elements taking part in the model and their connections. High concentration of the *driver* in the root tip confers cell proliferation activity. In turn, the *driver* promotes the expression of the cell division *regulator*. Cell cycle phase length is inversely proportional to regulator amounts inside the cell. Therefore, the model reproduces an incoherent feed forward loop system where the *driver* exerts a dual action on cell proliferation: it favors cell proliferation but at the same time it delays cell cycle progression. **F**, left panel: model simulation showing *driver* concentrations in the wild-type. High concentrations of a growth regulator in the root tip induces the expression of the *driver*, which is allowed to diffuse along the meristem. Middle panel: model simulation showing *regulator* concentrations in the wild-type. High *driver* concentrations in the root tip induces the expression of the *regulator*. Right panel: model simulation showing cell cycle time along the root as a result of *regulator* accumulation. **G**, Comparison between model simulation (light blue dots) and experimental data (red dots) in the wild-type. Each dot represents the data relative to a single cell. The *p-value* indicates the probability to obtain a similar data fit from random simulations (see Material and Methods). A *p-value* <0.05 indicates quantitative similarities between model predictions and experimental data. **H**, Comparison between model simulation (light blue dots) and experimental data (red dots) in the mutant with a reduction of the *driver* function (*plt1-4,plt2-2* mutant for experimental data). Statistical analysis as in panel G.

The AP2-ANT family caught our attention since it contains the AINTEGUMENTA-like (AIL)/PLETHORA (PLT) TFs, well known patterning genes with a gradient of expression in the RAM that control stem cell activity, confer cell proliferation potential and establish the longitudinal zonation in the RAM (Galinha *et al*, 2007; Mähönen *et al*, 2014; Aida *et al*, 2004). Therefore, PLT proteins fit with the features required to behave as a molecular driver of G1 length.

Single mutants of the four *PLT* genes do not exhibit a strong root phenotype, but *plt1-4,plt2-2* and *plt1-4,plt2-2,plt3-1* possess highly reduced root meristems with early termination, while higher order mutants exhibit rootless phenotypes (Galinha *et al*, 2007; Aida *et al*, 2004). Thus, we expressed the PlaCCI markers in the *plt1-4,plt2-2* double mutant background (Fig. 3B and video S2). RAM size was highly reduced in the *plt1-4,plt2-2* double mutant, as expected (Galinha *et al*, 2007; Mähönen *et al*, 2014; Aida *et al*, 2004). In addition, quantification of G1 duration by live imaging showed that the gradient observed in the wild type (Fig. 3C) was fully abolished in the *plt1-4,plt2-2* double mutant (Fig. 3D and Fig. S5). This supports a role for PLT proteins in the establishment of the G1 gradient in the root meristem.

Our mutant analysis showed that both *PLT*s affect negatively G1 progression, increasing G1 length, but the PLTs are also known to positively regulate cell proliferation potential (Galinha *et al*, 2007; Mähönen *et al*, 2014). Given the positive role of PLTs on meristem maintenance, we hypothesized the establishment of a G1 gradient could be achieved by the graded distribution of a positive regulator (e.g. PLTs) that induces a negative regulatory influence on G1. This regulatory relationship fits an incoherent feed-forward loop (IFFL) (Shen-Orr *et al*, 2002) entailing a common driver that regulates two branches with opposing effects on the output, one promoting cell proliferation potential and another promoting an asymmetric negative regulator that inhibits cell proliferation (Fig. 3E). To test whether an IFFL was sufficient to explain the G1 duration gradient, we adapted a computer model of the growing Arabidopsis root meristem that integrates the IFFL mechanism together with the spatial distribution of regulators and cell cycle characteristics (Marconi *et al*) (Supplementary Methods, Table S2). Here, the driver follows the graded distribution of a growth promoter (e.g., *PLT*s) and also controls the expression of a hypothetical negative G1 regulator. Spatial-temporal model simulations predict the pattern of cell cycle time along the RAM (Fig. 3F) and quantitatively matched the experimental observations of a proximal-distal gradient of G1 duration in the wild type. Removing the driver function of the IFFL in the model abolishes the G1 gradient compared with the wild type (Figs. 3G,H and Fig. S6). Therefore, our model shows that a PLT-initiated IFFL circuit is a plausible fold-change detector of PLT levels (Goentoro *et al*, 2009) for generating the robust G1 gradient along the meristem. However, our model also points to a gap in the regulatory circuit that would fill the role of the negative regulatory branch.

In a search for mechanisms that lie downstream of the PLTs, ChIP experiments have identified PLT2 target genes, including the CDK inhibitor KRP5/ICK3 (Santuari *et al*, 2016). We confirmed that the expression pattern of KRP5-GFP shows a gradient with high levels close to the QC that progressively decreased along the RAM (Fig. 4A and Fig. S7), consistent with the PLT gradient (Mähönen *et al*, 2014). It is worth noting that among the seven KRP family members, only the *krp5* mutants are significantly affected in primary root growth (Wen *et al*, 2013). Furthermore, the negative regulator pattern fit by the in-silico model closely matches the expression of KRP5-GFP expression in the meristem (Fig. S8). To determine the role of KRP5 in the control of G1 duration we expressed the PlaCCI markers in the *krp5-1* mutant background (Fig. 4B, left panel). We found that the *krp5-1* mutation led to a high number of cells in the distal half of the meristem that developed a faster G1, partially phenocopying the loss of *PLT* driver function (Fig. 4B, right panel, video S3 and Fig. S9).

**Fig. 4.**
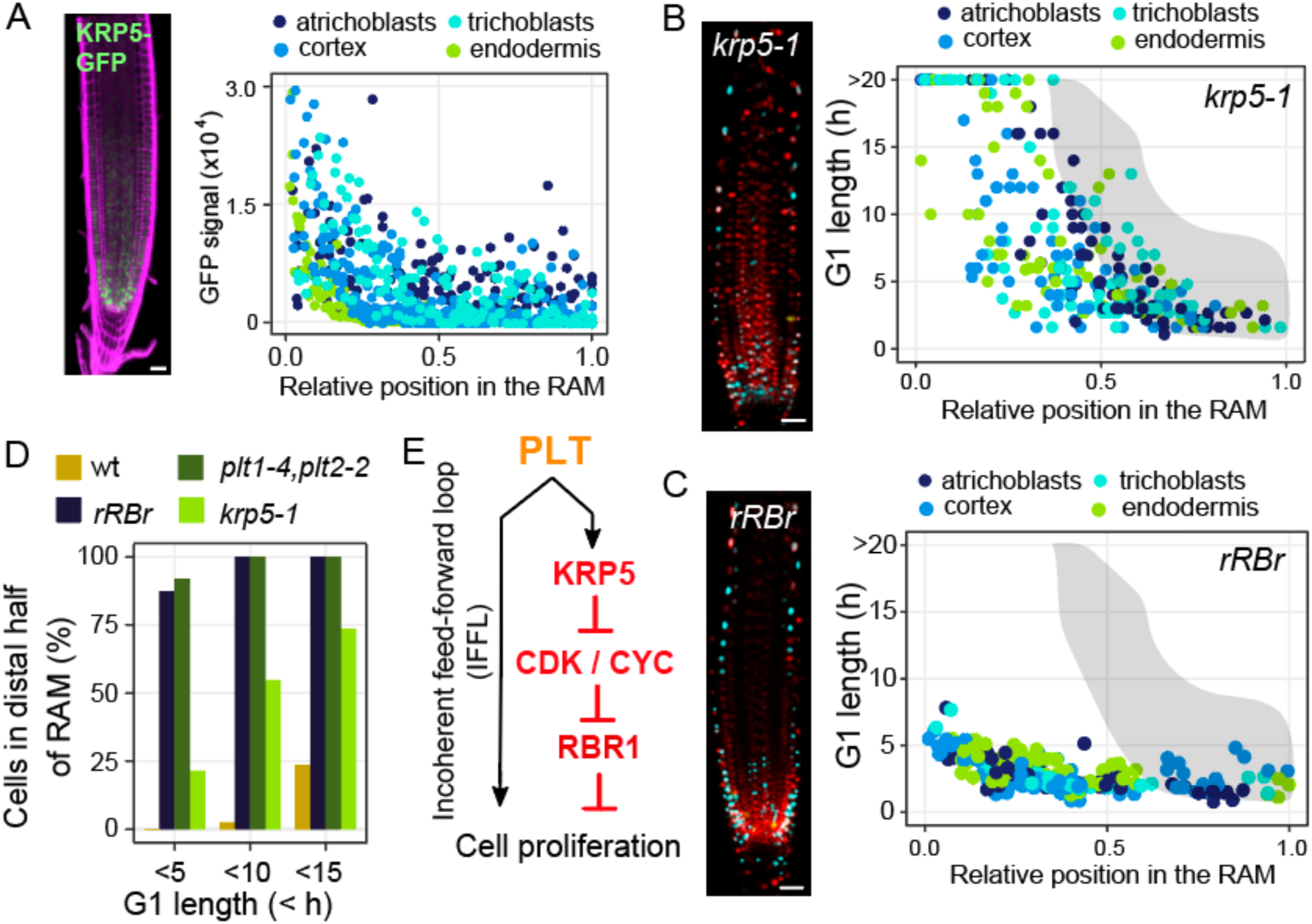
The RBR1-KRP5 pathway controls G1 duration and links with the *PLT* root patterning genes. **A**, Upper panel: root tip of a plant expressing the KRP5-GFP translational fusion expressed under the *KRP5* promoter. Scale bar = 20 µm. Bottom panel: quantification of the KRP5-GFP signal along the RAM (n = 791 cells). The RAM was divided into five sectors covering each 20% of the RAM, as indicated. 0 = QC, 1.0 = RAM boundary. **B**, Left panel: root tip of a *krp5-1* mutant expressing the PlaCCI markers. Scale bar = 20 µm. Right panel: G1 duration in four root cell types (atrichoblasts, trichoblasts, cortex, endodermis; n = 78, 73, 68 and 59, respectively), as indicated by the color code, along the root apical meristem (RAM) in the *krp5-1* mutant. The shaded shape is taken from Fig. 3C and is included here to facilitate comparison with the values of G1 duration in the wild type. **C**, Left panel: root tip of the *rbr1* loss-of-function mutant (*rRBr*) expressing the PlaCCI markers. Scale bar = 20 µm. Right panel: G1 duration in four root cell types (atrichoblasts, trichoblasts, cortex, endodermis; n= 62, 44, 94 and 95, respectively), as indicated by the color code, along the root apical meristem (RAM) in the the *rRBr* mutant. The shaded shape is taken from Fig. 3C and is included here to facilitate comparison with the values of G1 duration in the wild type. **D**, Quantification of the number of cells with different G1 durations in the distal half of the RAM in wild type and mutant roots. Data are taken from those shown in Figs. 3C, 3D, 4B and 4C. **E**, Basic components of the incoherent feed-forward loop (IFFL) identified in this study that controls the developmental regulation of G1 duration in the root apical meristem. PLT proteins that show a proximal-distal gradient in the RAM play an opposing function: they confer cell proliferation potential and also activates expression of the CDK inhibitor KRP5, which in turn restricts RBR1 phosphorylation and prolongs the G1 phase.

Since KRPs inhibit CDK activity, which in turn relieves from RBR1 repression, we focused on the RETINOBLASTOMA-RELATED 1 (RBR1) protein, which was shown to repress G1 progression (Desvoyes & Gutierrez, 2020). Thus, we followed G1 progression by live-imaging in RAM cells in *RBR1* loss-of-function mutant plants, using a viable *RBR1* RNAi line (*rRBr*) expressed specifically in RAM cells (Wildwater *et al*, 2005). Contrary to wild type meristems, where many CDT1a-CFP positive cells accumulate in the distal part of the RAM (Fig. 1A), the *rRBr* meristems showed significantly fewer CDT1a-CFP positive cells, indicating faster G1 progression than wild type in the distal meristem (Desvoyes *et al*, 2020) (Fig. 4C, left panel and video S4). Time-lapse experiments showed that 93.7% of cells in the whole RAM progress through G1 in less than ∼5 h in the absence of RBR1, fully abolishing the proximal-distal G1 length gradient observed in the wild type (Fig. 4C, right panel Fig. S10), consistent with RBR1 acting very downstream of the pathway. Thus, we conclude that RBR1 directly controls the developmentally regulated G1 duration along the proximal-distal axis of the root. Thus, RBR1 fulfills the role as part of the negative regulatory branch in the IFFL model, where the PLTs provide a driver mechanism.

An analysis of the G1 duration in the distal half of the meristem, which in the wild type contains only 2.5% of cells with a G1 <10h, revealed that this value was increased up to 54.7%, 100% and 100% in the *krp5-1, rRBr* and *plt1-4,plt2-2* mutants, respectively (Fig. 4D). This trend was increased further, showing 73.6% of cells having a G1 <15h in the *krp5-1* mutant compared to only 23.4% in the wild type (Fig. 4D). Our results indicate that KRP5 and RBR1 are part of the negative regulatory branch controlling the G1 length along the RAM.

The proposed gene regulatory network prevents unrestricted cell proliferation activity by the IFFL model with two antithetic branches: (i) one where PLTs stimulate cell proliferation potential, and (ii) another where PLTs activate *KRP5* expression, restricting cell cycle progression by lengthening the G1 phase. Other factors reported to control *PLT* gene expression such as XAL1/AGL12 (Tapia-Lopez *et al*, 2008; García-Cruz *et al*, 2016) or GRFs and miR396 (Ercoli *et al*, 2018; Liebsch & Palatnik, 2020) may cooperate in the control of G1 length along the RAM. The fold-change detection of PLT levels along the root provides a robust mechanism for controlling G1 duration and, hence, cell proliferation patterning during root development. In this scenario, the RBR1-KRP5 module works as the negative *regulator* in our IFFL model, where KRP5 is directly responsive of transferring the diffusion-mediated PLT gradient to the RBR1 regulatory function (Figure 4E). Our results uncover a new developmental feature of the longitudinal zonation of the root whereby a G1 duration gradient, dependent on the KRP5-RBR1 module, is directly linked with the activity of stem cell maintenance and patterning genes.

Our findings open up the investigation of the role of G1 length in plant development. In addition, the establishment of RBR1 in regulating a G1 gradient raises broader questions, as the role of retinoblastoma (Rb) proteins in G1 progression is highly conserved in eukaryotes (Desvoyes & Gutierrez, 2020). For example, in intestinal crypts, the Lgr5^+^-expressing stem cells develop longer cell cycles than their derivatives as a result of a longer G1 phase (Schepers *et al*, 2011; Carroll *et al*, 2018). A key question is whether fine-tuned control of G1 length by Rb-like proteins potentiates maturation gradients and developmental decisions in cell fate and potency.

## Materials and Methods

### Plant material and growth conditions

*Arabidopsis thaliana* lines used in this work are the following: wild type Columbia ecotype (Col-0) from NASC, PlaCCI (Desvoyes *et al*, 2020), *pRCH1::RBR* RNAi (rRBr) (Wildwater *et al*, 2005), *plt1-4,plt2-2* (Aida *et al*, 2004) and *krp5-1* (TAIR accession SALK_053533) (Wen *et al*, 2013). Plants expressing KRP5-GFP were generated by using a construct containing a 4363 insert of a genomic fragment spanning from 2989 bp upstream of the TSS up to 621 bp downstream of the stop codon. The PCR-amplified genomic fragment was cloned into the Gateway entry vector pDONR221 by BP reaction according to the manufacturer’s instructions (Thermo Fisher Scientific), and the sGFP tag was inserted just before the STOP codon by the SLiCE method (Motohashi, 2015). An LR reaction was performed with the destination vector pGWB1 (Nakagawa *et al*, 2007) to generate a binary vector. Seeds were bleach-sterilized, stratified at 4 °C for 2 days and grown in agar plates containing 0.5x Murashige-Skoog (MS) medium (pH 5.7) supplemented with MES (Sigma), vitamins, 1% sucrose, and 0.6, 0.8 or 1% agar (Duchefa). Plant transformation was carried by the floral dip method (Clough & Bent, 1998). Plants grew vertically or horizontally in an incubator at 20 °C and 60% moisture, under long-day conditions (16 h light/8 h dark cycles, fluorescent tubes Radium Spectralux Plus NL-T8, 36 W, cool daylight, 100 μmol m-2 s-1).

### In vivo imaging and confocal microscopy

Seedlings were transferred 4 days post-sowing (dps) to P35 coverslip bottom dishes (MatTek). A block of 1% agar 0.5xMS medium was placed on top of the seedlings, cutting a segment with a sterilized blade, making room for the aerial part to grow outside of the agar. To avoid curling and drifting of the roots in the z-plane, a square piece of 100 µm nylon mesh (Nitex) was placed between the seedlings and the agar block, creating tracks for the roots to grow straight. After fixing the lid with Micropore tape, roots were allowed to acclimate overnight, placing dishes vertically in the plant culture chamber. Root meristems were imaged every 30 minutes up to 20 hours with a Nikon A1R+ inverted confocal microscope using an oil 40x objective. Confocal stacks spanning over half the root thickness (pinhole of 3.5-4 µm, ∼50 µm distributed in ∼14 Z-planes) were acquired at each time point, with 2×1 tile-scan to cover the whole meristem plus the endoreplication zone length. Images and video editing were performed using FIJI (Rueden *et al*, 2017). Different root cell types were identified by their anatomical position in the root. In each case, scoring was initiated at the RAM boundary and then continuing along the file up to the QC.

Positioning of cells and mitotic figures in the root meristem

The position of individual cells in the meristem was calculated by:

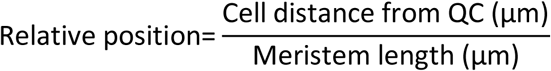

Meristem length was taken as the distance from the QC to the first cell doubling its size. Thus, a relative position of 0 corresponds to the QC and 1 to the end of the meristem. Epidermal, cortical and endodermal cell positions were relative to their own tissue meristems. When this position was calculated in the analysis of in vivo experiments, the cell distance from the QC was that at the beginning of the process under study, for example, right after the anaphase of a cell in which G1 was being measured.

### EdU labeling and chase experiments

Seedlings (7 days after sowing) were transferred to 0.5x liquid MSS containing 20 µM EdU for 15 min to label cells undergoing S-phase. Then the analogue was washed off and plants were incubated in 0.5x liquid MSS supplemented with 50 µM thymidine (Sigma) for different chase time periods to allow cell cycle progression. Plants were then fixed with 4% paraformaldehyde in microtubule stabilizing buffer (MTSB; 50 mM PIPES, pH 6.9, 5 mM EGTA, 5 mM MgSO4) and permeabilized as described (Lauber *et al*, 1997). EdU incorporation was detected with the Click-it Alexa Fluor 647 Imaging kit (Life Technologies), nuclei stained with DAPI, and roots imaged by confocal microscopy with a Zeiss LSM800. G2 phase length was directly measured along the root meristem by determining the kinetics of appearance of EdU-labeled mitotic figures in the different tissues and chase times (Fiorani & Beemster, 2006).

### Analysis of the dynamics of cell cycle marker proteins

Residence time of CYCB1;1-YFP in the cells was measured by in vivo imaging of PlaCCI seedlings (Desvoyes *et al*, 2020), as the time between the loading of this protein in G2 and its degradation at the metaphase-anaphase transition. CDT1a loading kinetics was assessed by measuring the time between the end of anaphase and a detectable CDT1a-CFP signal. Likewise, G1 phase duration was given by the time between the end of mitosis and the degradation of CDT1a-CFP at the G1-S transition.

### Differential expression of transcription factors along the RAM

Data were taken from the spatial gene expression patterns along the root obtained from correlative root slices from the tip (slice 1) to the basal part of the root (Brady). We first selected a set of 1,472 genes with variable expression along the root and clustered them using weighted correlation network analysis (WGCNA) to identify modules of highly correlated genes. We obtained 7 modules (M1 through M7) containing genes of highly correlated expression levels. These were classified into three groups depending of their profiles as type A, B and C modules, corresponding to the highest expression in distal (1-6), middle (6-7) or proximal slices (10-12), respectively. To identify transcription factors (TFs) putatively contributing to the establishment of the expression profiles we searched for enriched transcription factor binding sites (TFBS) in the promoters of genes in each module. TFBS were mapped in 1 kb gene promoters with FIMO, and fold-enrichment in log2 scale calculated by mapping the same sites in the complete Arabidopsis promoter set. A heat map was calculated for individual TF families identified.

### Computational model

The model was built using MorphoDynamX, a modeling platform based on MorphoGraphX (Barbier de Reuille *et al*, 2015). MorphoDynamX is an advanced modeling framework based on a geometric data structure called Cell Complexes (Lane, 2015). The mechanical growth of the simulated root is based on Position-Based Dynamics (PBD), a modern constraint-based method used to simulate physical phenomena like cloth, deformation, fluids, fractures, rigidness and much more (Tsai, 2017). Chemical processes are numerically solved using the simple Euler method. The current model is an extension of a previous version, and we refer the reader to the original publication for details about the model implementation (Marconi, M. *et al*). The presented version expands the previous one by adding chemical control over cell division, according to the observations described in the main text. Cell division is regulated by two chemical components, a *driver* and a *regulator*. The presence of the division *driver* allows the cell to divide, while the *regulator* negatively regulates the cell division cycle. Higher concentrations of the *regulator* correlate with longer division cycles. The system follows the scheme described by an incoherent feed-forward loop (IFFL; Fig. 3E). The full list of parameters description can be found in Supplementary Table S2. Specifically, the division *driver* is induced by an upstream growth promoter, which is assumed to be present at high concentrations in the root tip. The *driver* is allowed to diffuse between cells (Mähönen *et al*, 2014), according to the formula:

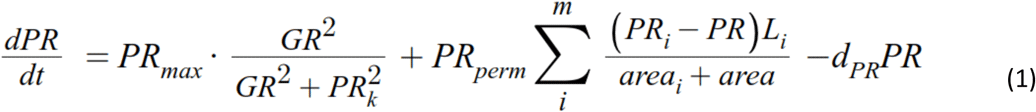

*PR*_*max*_ is the maximum level of growth promoter-induced *driver* expression; *PR*_*k*_ is the half-max coefficient of growth promoter-induced *driver* function; *PR*_*perm*_ is the *driver* cell permeability coefficient; *d*_*PR*_ is the *driver* degradation rate; *GR* is the growth promoter concentration inside the current cell; *PR* is the *driver* concentration inside the current cell; *area* is the total area of the current cell; the summation symbol indicates iteration over the (*m*) neighbor cells of the current cell, while *PR*_*i*_ and *area*_*i*_ indicate *driver* concentration and area of a specific neighboring cell, respectively. *L*_*i*_ is the length of the membrane between the current cell and the specific neighboring cell.

Conversely, *regulator* expression is promoted by the *driver*:

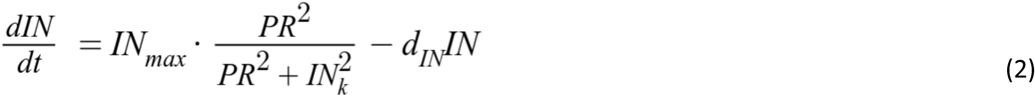

*IN*_*max*_ is the maximum level of *driver*-induced *regulator* function; *IN*_*k*_ is the half-max coefficient of *driver*-induced *regulator* function; *d*_*IN*_ is the *regulator* degradation rate; *PR* is the *driver* concentration inside the current cell.

The cell cycle time (the minimum amount of time between cell divisions, which is considered here directly proportional to the G1 length) is directly regulated by the *regulator* according to the following formula:

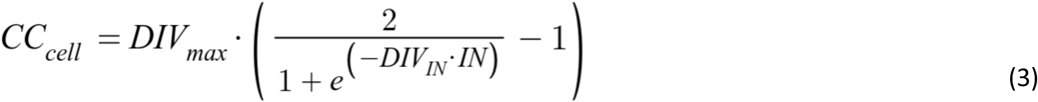

*DIV*_*max*_ is the maximum cell cycle time; *DIV*_*IN*_ is the exponential rate of cell cycle regulation by the *regulator*; *IN* is the concentration of the *regulator* inside the current cell.

Considering the aforementioned components, cell division is implemented in the simulations according to the following rules:

- Only cells inside the meristem can divide.
- Cells are allowed to divide only once they have doubled their size in length.
- The division *driver* allows cell division once its concentration is above a certain threshold, *DIV*_*prom*_.
- A cell can divide only if its lifetime (the time passed since the last division) has elapsed the cell cycle time determined by the division *regulator, CC*_*cell*_.
- The division *regulator* is degraded after cell division.

A statistical analysis has been performed to obtain hypothesis testing results for the experimental data. We calculate the Hausdorff distance (Birsan & Tiba, 2006) between experimental data and model simulations for both the wild type and mutant. Next, we generated 10000 random pairs of data from the following formula:

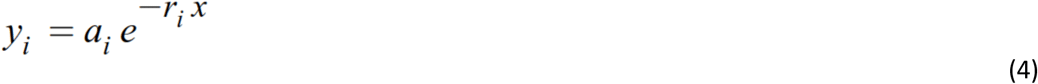

Where *i* ∈[1,2] indicates the pair element index, *a* ∈(0, 50] indicates the curve intercept (in accordance to the maximum cell cycle length), and r ∈(0,10] indicates the exponential disgrace in cell cycle length (in accordance with the observed experimental data). For each of the previous random pairs we calculated the Hausdorff distance, and finally calculated the *p-value* as the proportion of random distances smaller than the Hausdorff distance obtained between experimental data and model simulations. A small *p-value* indicated that the observed similarity between experimental data and model simulations is hard to explain by chance alone.

## Acknowledgments

We thank B. Scheres and R. Heidstra for the *rRBr* RNA line and the *plt1-4,plt2-2* seeds, J. Murray for the *krp5-1* mutant, the Confocal Microscopy Service of CBM, D. Miguez for initial discussions on root modeling and E. Martinez-Salas, M. Serrano and members of our laboratories for comments and suggestions.

## Author contributions

Conceptualization: CG, BD, KW

Methodology: CE, BD, MM, LL, JMF-Z, RS, MU, KB

Investigation: CE, BD, MM, LL, JMF-Z

Funding acquisition: CG, KW, JMF-Z, RS, KB

Project administration: CG, BD Supervision: CG, BD, KW, RS, KB

Writing – original draft: CG, BD

Writing – review & editing: all authors

## Data and materials availability

All data are available in the main text or the supplementary materials.

## Supplementary Materials

Figs. S1 to S10

Tables S1 and S2

References

Movies S1 to S4

## References

Aida M, Beis D, Heidstra R, Willemsen V, Blilou I, Galinha C, Nussaume L, Noh YS, Amasino R & Scheres B (2004) The PLETHORA genes mediate patterning of the Arabidopsis root stem cell niche. Cell 119: 109–20

Barbier de Reuille P, Routier-Kierzkowska A-L, Kierzkowski D, Bassel GW, Schüpbach T, Tauriello G, Bajpai N, Strauss S, Weber A, Kiss A, et al (2015) MorphoGraphX: A platform for quantifying morphogenesis in 4D. Elife 4: 05864

Birsan, T. & Tiba, D. (2006) One Hundred Years Since the Introduction of the Set Distance by Dimitrie Pompeiu. In System Modeling and Optimization. CSMO 2005. IFIP International Federation for Information Processing Springer, Boston, MA

Boward B, Wu T & Dalton S (2016) Concise Review: Control of Cell Fate Through Cell Cycle and Pluripotency Networks. Stem Cells 34: 1427–1436

Brady SM, Orlando DA, Lee JY, Wang JY, Koch J, Dinneny JR, Mace D, Ohler U & Benfey PN (2007) A high-resolution root spatiotemporal map reveals dominant expression patterns. Science 318: 801–806

Carroll TD, Newton IP, Chen Y, Blow JJ & Näthke I (2018) Lgr5(+) intestinal stem cells reside in an unlicensed G(1) phase. J Cell Biol 217: 1667–1685

Clough SJ & Bent AF (1998) Floral dip: a simplified method for Agrobacterium-mediated transformation of Arabidopsis thaliana. Plant J 16: 735–743

Clowes, F.A.L. (1965) The duration of the G1 phase of the mitotic cycle and its relation to radiosensitivity. New Phytol 64: 355–359

Costa S & Shaw P (2006) Chromatin organization and cell fate switch respond to positional information in Arabidopsis. Nature 439: 493–496

Desvoyes B, Arana-Echarri A, Barea MD & Gutierrez C (2020) A comprehensive fluorescent sensor for spatiotemporal cell cycle analysis in Arabidopsis. Nat Plants 6: 1330–1334

Desvoyes B & Gutierrez C (2020) Roles of plant retinoblastoma protein: cell cycle and beyond. EMBO J 39: e105802

Dolan L, Janmaat K, Willemsen V, Linstead P, Poethig S, Roberts K & Scheres B (1993) Cellular organisation of the Arabidopsis thaliana root. Development 119: 71–84

Echevarría, C., Gutierrez, C. & Desvoyes, B. (2021) Tools for Assessing Cell Cycle Progression in Plants. Plant Cell Physiol 62: 1231–1238

Ercoli MF, Ferela A, Debernardi JM, Perrone AP, Rodriguez RE & Palatnik JF (2018) GIF Transcriptional Coregulators Control Root Meristem Homeostasis. Plant Cell 30: 347–359

Fiorani F & Beemster GTS (2006) Quantitative analyses of cell division in plants. Plant Mol Biol 60: 963–979

Galinha C, Hofhuis H, Luijten M, Willemsen V, Blilou I, Heidstra R & Scheres B (2007) PLETHORA proteins as dose-dependent master regulators of Arabidopsis root development. Nature 449: 1053–1057

García-Cruz KV, García-Ponce B, Garay-Arroyo A, Sanchez Mdlp, Ugartechea-Chirino Y, Desvoyes B, Pacheco-Escobedo MA, Tapia-López R, Ransom-Rodríguez I, Gutierrez C, et al (2016) The MADS-box XAANTAL1 increases proliferation at the Arabidopsis root stem-cell niche and participates in transition to differentiation by regulating cell-cycle components. Ann Bot 118: 787–796

Goentoro L, Shoval O, Kirschner MW & Alon U (2009) The incoherent feedforward loop can provide fold-change detection in gene regulation. Mol Cell 36: 894–899

Ivanov VB & Dubrovsky JG (1997) Estimation of the cell-cycle duration in the root meristem: a model of linkage between cell-cycle duration, rate of cell production, and rate of root growth. Int J Plant Sci 158: 757–763

Lane BJ (2015) Cell Complexes: The Structure of Space and the Mathematics of Modularity. doi:10.11575/PRISM/10182 [PREPRINT]

Lauber MH, Waizenegger I, Steinmann T, Schwarz H, Mayer U, Hwang I, Lukowitz W & Jurgens G (1997) The Arabidopsis KNOLLE protein is a cytokinesis-specific syntaxin. J Cell Biol 139: 1485–1493

Liebsch D & Palatnik JF (2020) MicroRNA miR396, GRF transcription factors and GIF co-regulators: a conserved plant growth regulatory module with potential for breeding and biotechnology. Curr Opin Plant Biol 53: 31–42

Mähönen AP, ten Tusscher K, Siligato R, Smetana O, Diaz-Trivino S, Salojarvi J, Wachsman G, Prasad K, Heidstra R & Scheres B (2014) PLETHORA gradient formation mechanism separates auxin responses. Nature 515: 125–129

Marconi, M., Gallemi, M., Benkova, E., & Wabnik, K. A coupled mechano-biochemical framework for root meristem morphogenesis. BioRxiv https://doi.org/10.1101/2021.01.27.428294

Meyer HM, Teles J, Formosa-Jordan P, Refahi Y, San-Bento R, Ingram G, Jönsson H, Locke JCW & Roeder AHK (2017) Fluctuations of the transcription factor ATML1 generate the pattern of giant cells in the Arabidopsis sepal. Elife 6:e19131

Motohashi K (2015) A simple and efficient seamless DNA cloning method using SLiCE from Escherichia coli laboratory strains and its application to SLiP site-directed mutagenesis. BMC Biotechnol 15: 47

Nakagawa T, Kurose T, Hino T, Tanaka K, Kawamukai M, Niwa Y, Toyooka K, Matsuoka K, Jinbo T & Kimura T (2007) Development of series of gateway binary vectors, pGWBs, for realizing efficient construction of fusion genes for plant transformation. J Biosci Bioeng 104: 34–41

Otero S, Desvoyes B, Peiro R & Gutierrez C (2016) Histone H3 dynamics uncovers domains with distinct proliferation potential in the Arabidopsis root. Plant Cell 28: 1361–1371

Pacheco-Escobedo MA, Ivanov VB, Ransom.Rodriguez I, Arriaga-Mejía G, Avila H, Baklanov IA, Pimentel A, Corkidi G, Doerner P, Dubrovsky JG, et al (2016) Longitudinal zonation pattern in Arabidopsis root tip defined by multiple structural change algorithm. Ann Bot 118: 763–776

Pauklin S & Vallier L (2013) The cell-cycle state of stem cells determines cell fate propensity. Cell 155: 135–147

Rahni R & Birnbaum KD (2019) Week-long imaging of cell divisions in the Arabidopsis root meristem. Plant Methods 15: 30

Rahni R, Efroni I & Birnbaum KD (2016) A Case for Distributed Control of Local Stem Cell Behavior in Plants. Dev Cell 38: 635–642

Rueden CT, Schindelin J, Hiner MC, DeZonia BE, Walter AE, Arena ET & Eliceiri KW (2017) ImageJ2: ImageJ for the next generation of scientific image data. BMC Bioinformatics 18: 529

Sablowski, R. & Gutierrez, C. (2021) Cycling in a crowd: coordination of plant cell division, growth and cell fate. Plant Cell: 193–208

Santuari L, Sanchez-Perez GF, Luijten M, Rutjens B, Terpstra I, Berke L, Gorte M, Prasad K, Bao D, Timmermans-Hereijgers JLPM, et al (2016) The PLETHORA Gene Regulatory Network Guides Growth and Cell Differentiation in Arabidopsis Roots. Plant Cell 28: 2937–2951

Schepers AG, Vries R, van den Born M, van de Wetering M & Clevers H (2011) Lgr5 intestinal stem cells have high telomerase activity and randomly segregate their chromosomes. EMBO J 30: 1104–1109

Scheres B (2007) Stem-cell niches: nursery rhymes across kingdoms. Nature reviews 8: 345– 54

Shen-Orr SS, Milo R, Mangan S & Alon U (2002) Network motifs in the transcriptional regulation network of Escherichia coli. Nat Genet 31: 64–68

Tapia-Lopez R, Garcia-Ponce B, Dubrovsky JG, Garay-Arroyo A, Perez-Ruiz RV, Kim SH, Acevedo F, Pelaz S & Alvarez-Buylla ER (2008) An AGAMOUS-related MADS-box gene, XAL1 (AGL12), regulates root meristem cell proliferation and flowering transition in Arabidopsis. Plant Physiol 146: 1182–1192

Tsai T-C (2017) Position Based Dynamics. In Encyclopedia of Computer Graphics and Games, Lee N (ed) pp 1–5. Cham: Springer International Publishing

Wen B, Nieuwland J & Murray JAH (2013) The Arabidopsis CDK inhibitor ICK3/KRP5 is rate limiting for primary root growth and promotes growth through cell elongation and endoreduplication. J Exp Bot 64: 1135–1144

Wildwater M, Campilho A, Perez-Perez JM, Heidstra R, Blilou I, Korthout H, Chatterjee J, Mariconti L, Gruissem W & Scheres B (2005) The RETINOBLASTOMA-RELATED gene regulates stem cell maintenance in Arabidopsis roots. Cell 123: 1337–49

Zhukovskaya NV, Bystrova EI, Dubrovsky JG & Ivanov VB (2018) Global analysis of an exponential model of cell proliferation for estimation of cell cycle duration in the root apical meristem of angiosperms. Ann Bot 122: 811–822

