## Supplementary Material for "Stem cell regulators control a G1 duration gradient in the plant root meristem"

**This PDF file include:**

- Figs. S1 to S10
- Tables S1 to S2
- Captions for Movies S1 to S4
- References

**Other Supplementary Materials for this manuscript include the following:**

- Movies S1 to S4

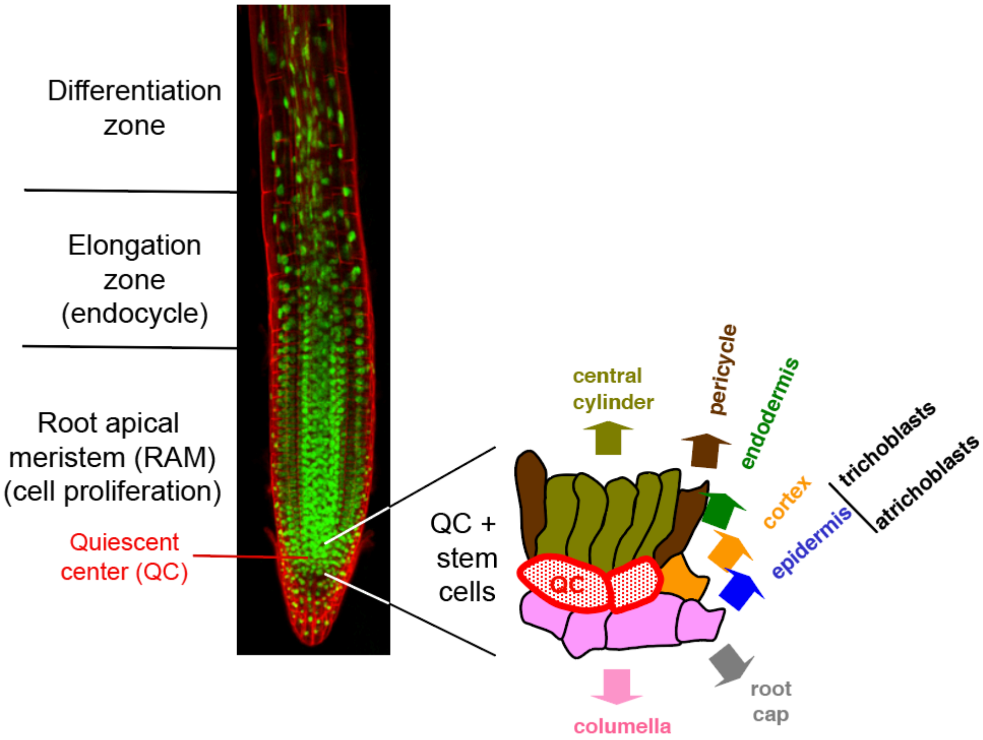


**Figure S1. Anatomical organization of the Arabidopsis root apex.** Nuclei are identified by the constitutive expression of a histone H3.3-GFP protein (green) and the cell walls were stained with propidium iodide. On the radial axis, concentrical cell layers can be identified from the outermost epidermal layer to the cortex, endodermis, pericycle and vascular tissues. On the longitudinal axis, the root apical meristem (RAM), the focus of our study, occupies the more distal (or rootward) zone and contains all proliferating cells (Desvoyes, B. *et al*, 2021). The RAM is followed by an elongtion zone, where cells stop proliferation, enter the endocycle and initiate elongation in the longitudinal axis, and finally a differentiation zone where cells acquire their final differentiated state. Within the RAM, a group of rarely dividing cells, the quiescent center (QC) cells, is located distally and is surrounded by stem cells (Scheres, 2007; Shimotohno & Scheres, 2019) that give rise to all different root cells types (see inset). Asymmetric division of these stem cells renders the first derivatives that undergo several divisions within the transit amplifying compartment before arresting the cell cycle and initiating an elongation process, coinciding with the RAM boundary (Pacheco-Escobedo *et al*, 2016), which is defined by a complex hormonal balance (Salvi *et al*, 2020; Svolacchia *et al*, 2020).

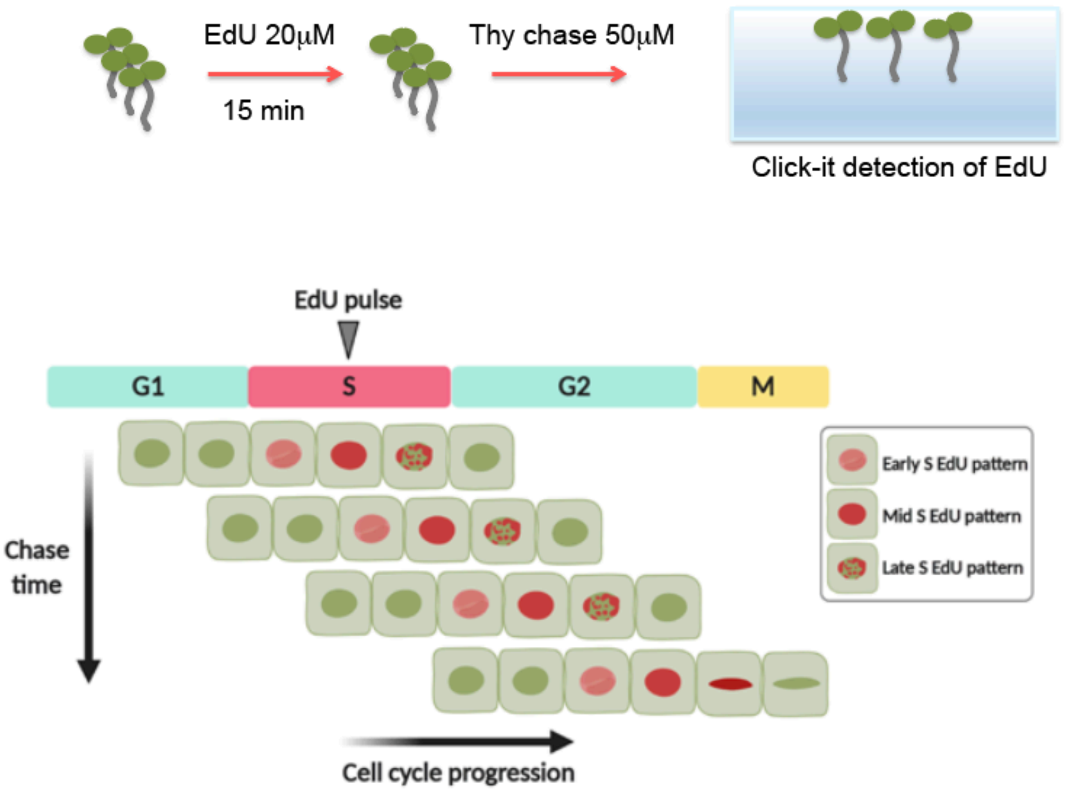


**Figure S2. EdU pulse-chase strategy for direct measurement of the G2 phase duration.** Seedlings were labeled with a short (15 min) pulse of the thymidine analogue EdU and then chased for different periods of time, in the absence of EdU and in the presence of an excess of thymidine to reduce further EdU incorporation, before Click-It detection of EdU labeled nuclei, as indicated in the upper part of the figure. The lower part of the figure contains a simplified scheme to illustrate cell cycle progression during the chase period. Cells in S-phase are all labeled with EdU during the pulse. Then, with time, cells progress across G2 and eventually enter mitosis. Quantification of mitotic cells labeled with EdU in the different chase times measures the G2 length. The average G2 phase duration corresponds to the chase time where 50% of mitotic cells appear labeled, assuming a normal distribution of the EdU labeled cells. This figure has been created with BioRender.com.

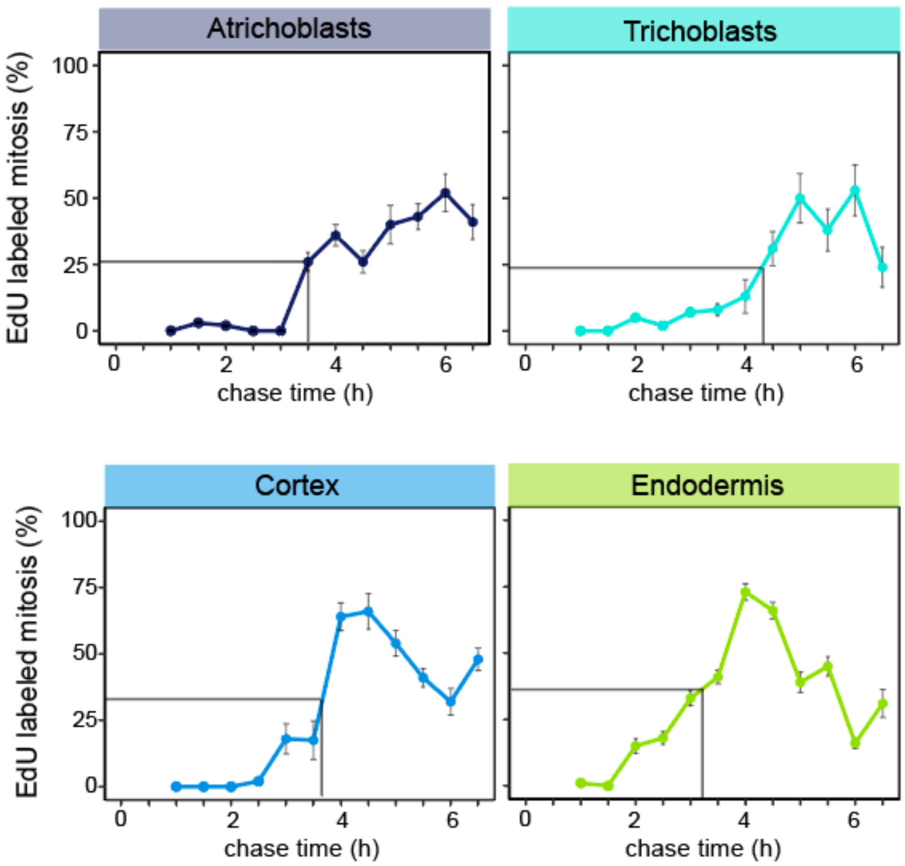


**Figure S3. Average duration of G2 in various cell types.** The fraction of EdU-labeled mitosis was quantified at different chase times after a pulse with EdU (15 min). The average G2 duration was estimated at 50% of the maximum reached in the curve (indicated by the thin lines in each case).

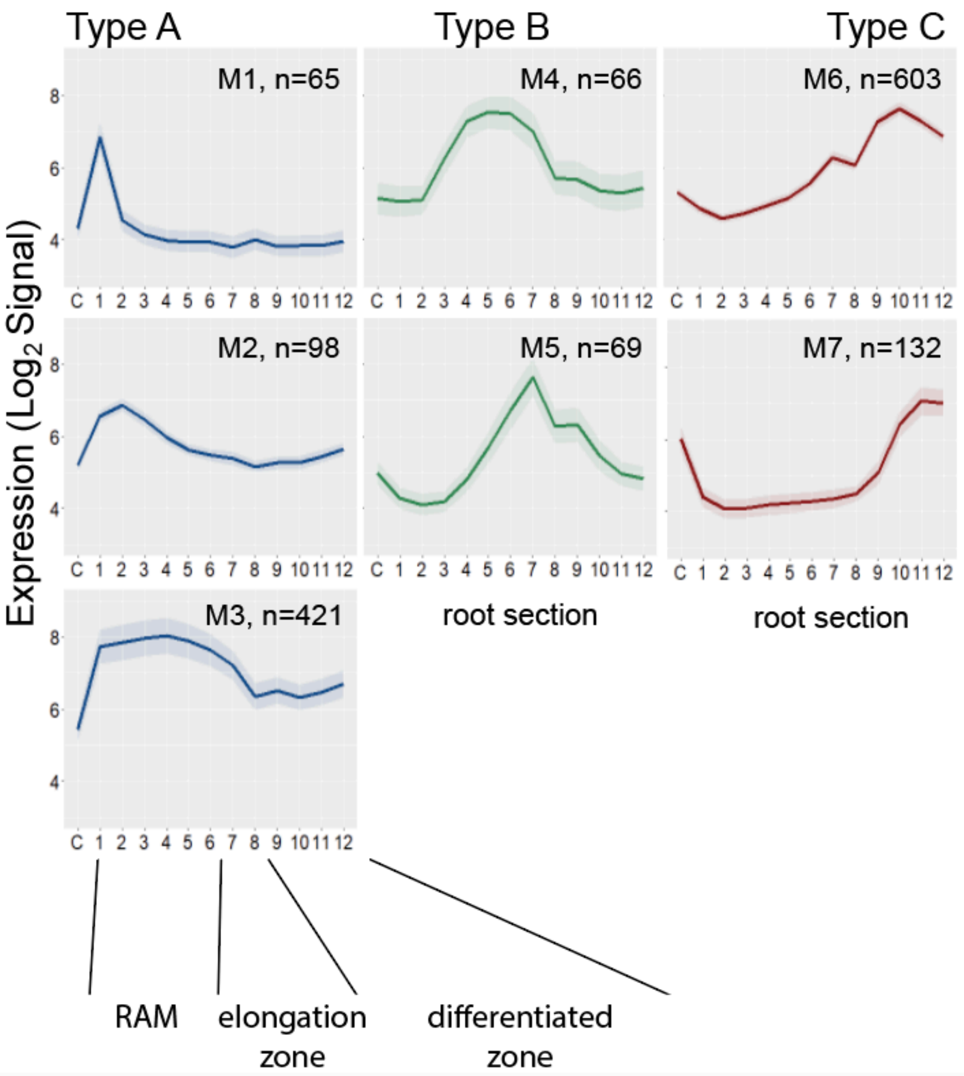


**Figure S4. WGCNA clustering in relation to gene expression along the RAM.** Root sections were taken from Brady et al. (Brady *et al*, 2007) where C corresponds to the columella and the RAM boundary to section 6. Plots represent mean gene expression (and confidence interval in shade) of the genes included in each of the modules or clusters. The number of genes is indicated as 'n'. Gene modules are classified in relation to their maxima expression in proximal (type A), middle (type B) and distal (type C) root sections (Section C = columella; sections 1-6 = RAM; sections 7-8 = transition (endoreplication) zone; sections 9-12 = elongation-differentiation zone).

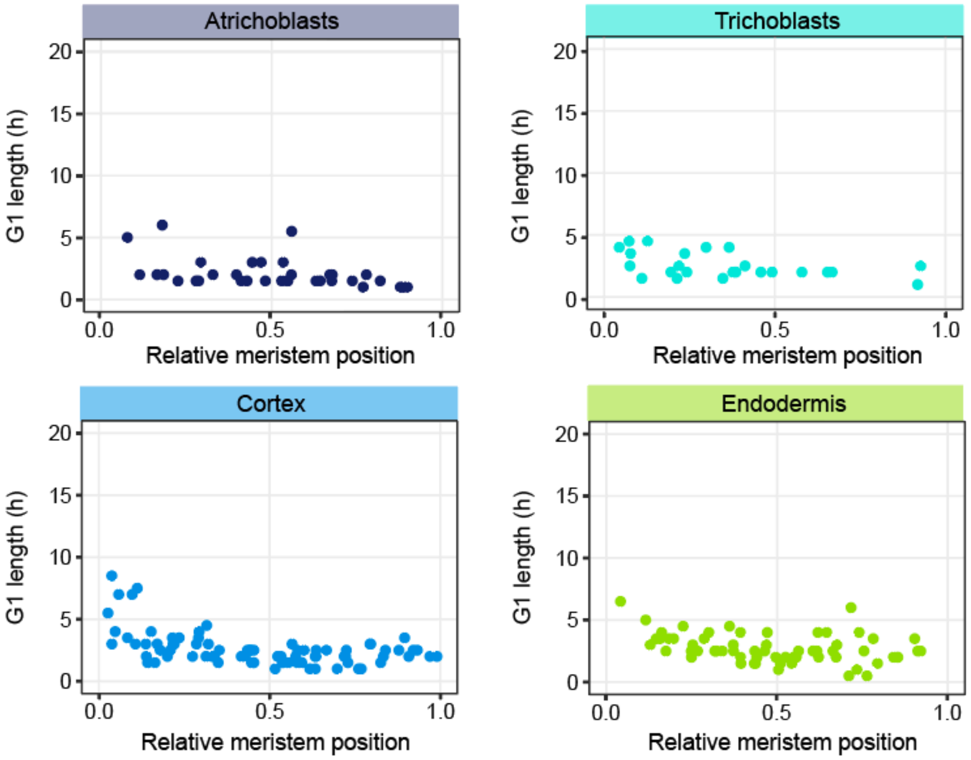


**Figure S5. Duration of G1 in various cell types in cells of *plt11-4,plt2-2* double mutant.** Videos were recorded covering the RAM of plants expressing the PlaCCI markers in a *plt11-4,plt2-2* double mutant, as described in the main text. Each dot represents the G1 length of individual cells. Cell numbers were 36, 26, 81 and 64 for atrichoblasts, trichoblasts, cortex and endodermis, respectively.

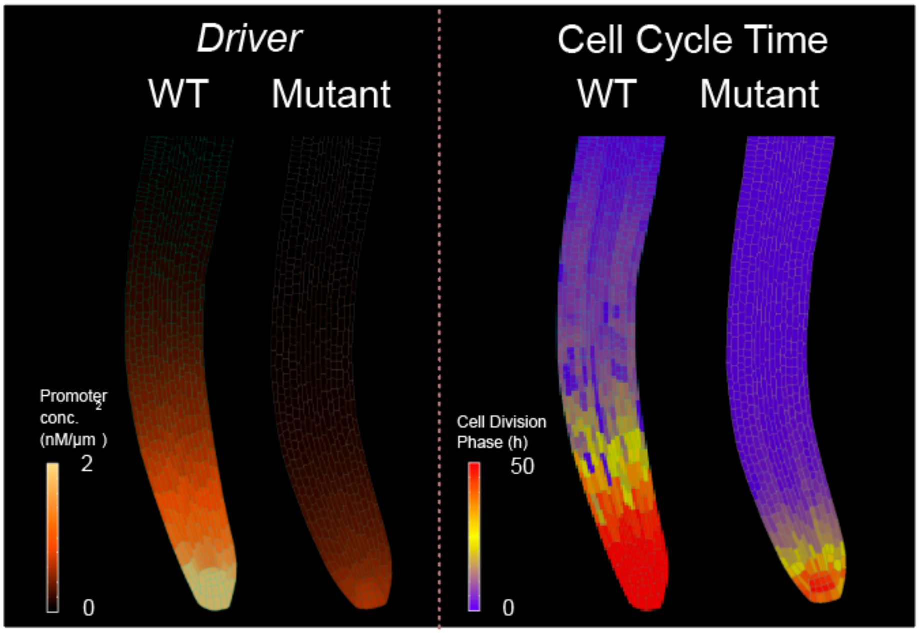


**Figure S6.** Left panel: model simulation showing *driver* activity in the wild type and *plt1-4,plt2-2* mutant. Right panel: model simulation showing cell cycle time (a proxy of G1 duration in our model) along the root as a result of *driver* accumulation in the wild type and the mutant.

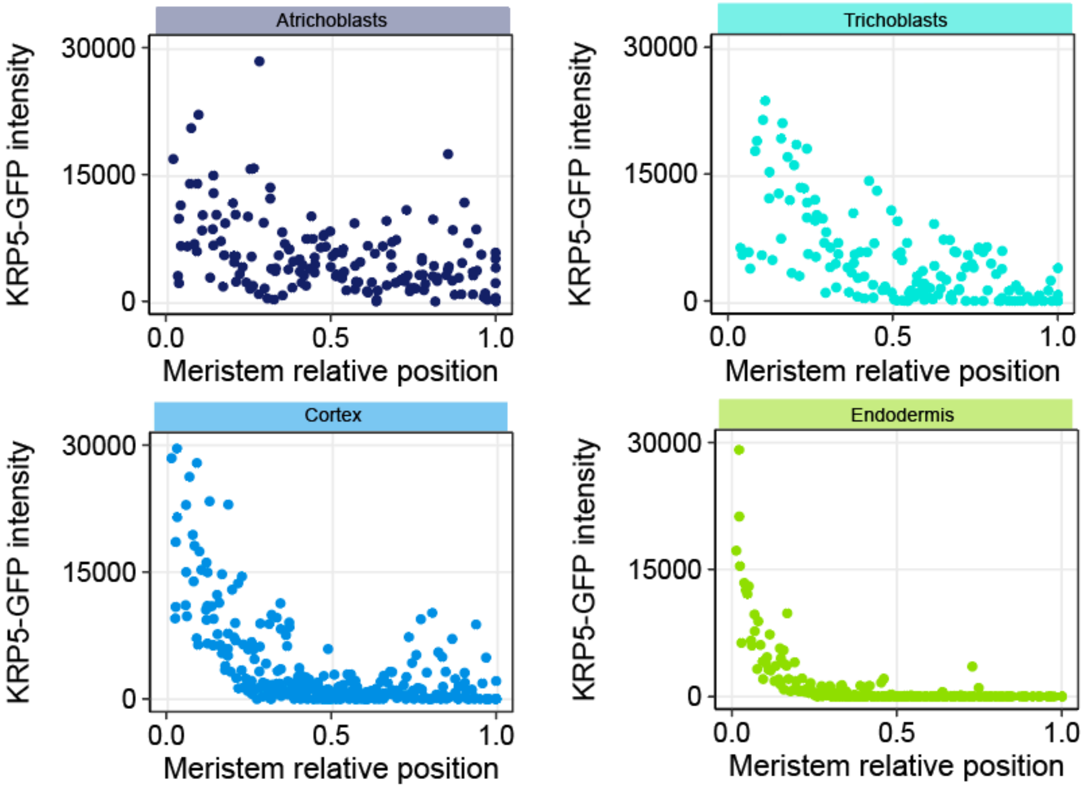


**Figure S7. Expression pattern of KRP5-GFP along the RAM in various cell types.** The GFP signal was quantified in individual cells and then grouped in classes spanning 0.2 units of the RAM. Cell numbers were 158, 146, 268 and 219 for atrichoblasts, trichoblasts, cortex and endodermis, respectively.

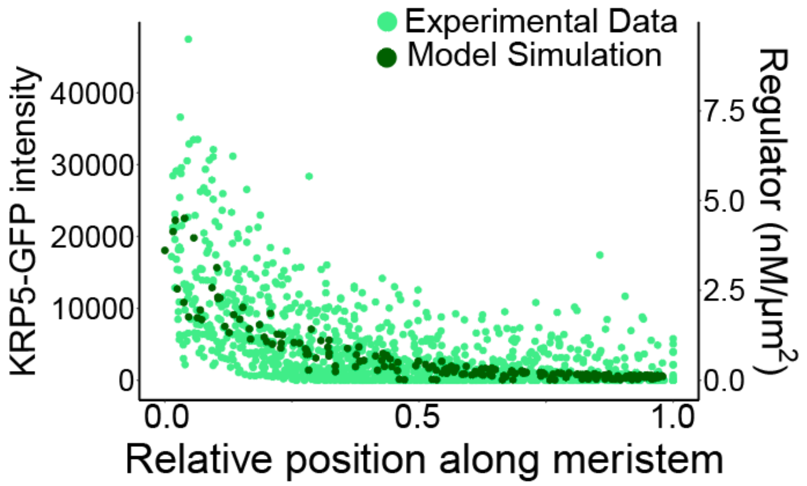


**Figure S8. Comparison between model simulation and experimental data of the KRP5-GFP signal.** Experimental KRP5-GFP data (light green dots) with model simulation data (dark green dots). Each dot represents the value of individual cells.

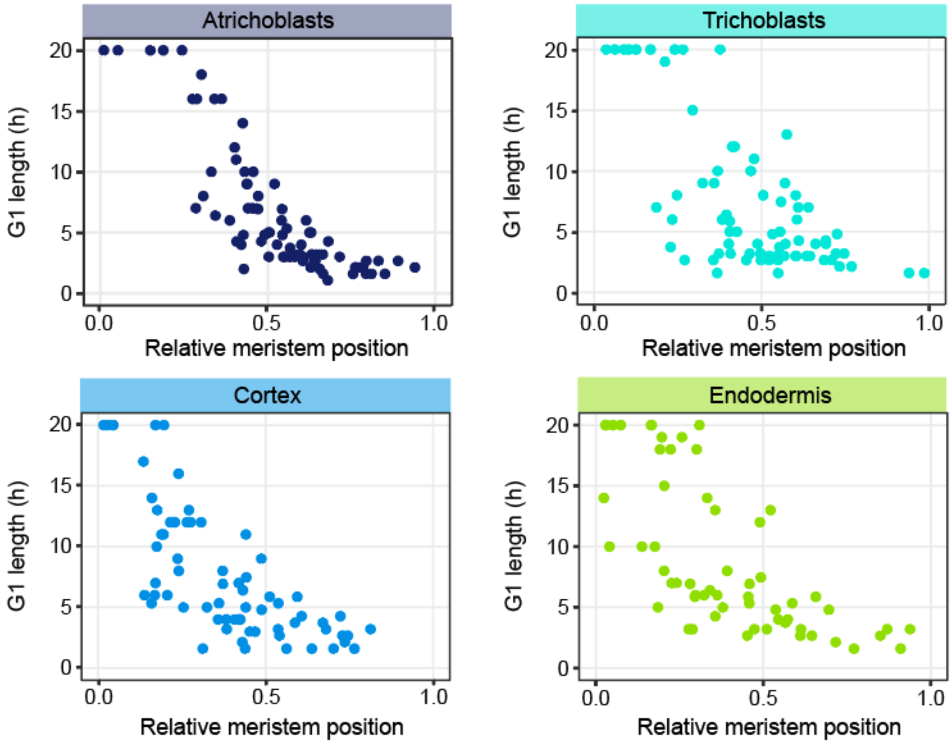


**Figure S9. Duration of G1 in various cell types in cells of *krp5-1* mutant.** Videos were recorded covering the RAM of plants expressing the PlaCCI markers in the *krp5-1* mutant, as described in the main text. Each dot represents the G1 length of individual cells. Cell numbers were 36, 26, 81 and 64 for atrichoblasts, trichoblasts, cortex and endodermis, respectively.

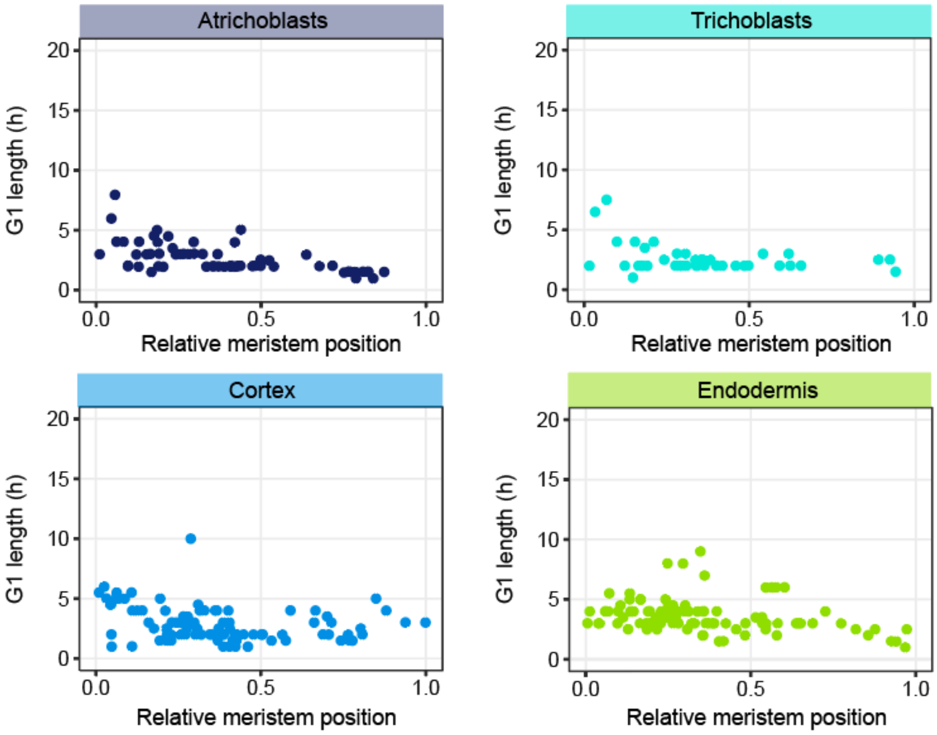


**Figure S10. Duration of G1 in various cell types in cells lacking the RBR1 protein.** Videos were recorded covering the RAM of plants expressing the PlaCCI markers in a mutant lacking RBR1 (*rBRr* plants), as described in the main text. Each dot represents the G1 length of individual cells. Cell numbers were 62, 44, 94 and 95 for atrichoblasts, trichoblasts, cortex and endodermis, respectively.

**Table S1. Clustering datasets of transcription factor binding sites in genes with high expression in the distal half of the RAM.**

This is provided as a separate excel file

**Table S2. Model parameters description and values.**

| Parameter | Description | Value | Units |
|---|---|---|---|
| $PR_{max}$ | maximum level of auxin-induced *driver* expression | 20 | nM/h |
| $PR_k$ | half-max coefficient of auxin-induced *driver* expressi | 2 | nM/$\mu m^2$ |
| $PR_{perm}$ | *driver* cell permeability coefficient | 1 | µm/h |
| $d_{PR}$ | *driver* degradation rate | 0.01 | $h^{-1}$ |
| $IN_{max}$ | maximum level of *driver*-induced *regulator* expressio | 20 | nM/h |
| $IN_k$ | half-max coefficient of *driver*-induced *regula* expression | 5 | nM/$\mu m^2$ |
| $d_{IN}$ | *regulator* degradation rate | 0.01 | µm/h |
| $DIV_{max}$ | maximum cell cycle time | 50 | h |
| $DIV_{IN}$ | exponential rate of cell cycle regulation by the *regula* | 0.03 | $h^{-1}$ |

For a global description of the default parameters please refer to the original publication (Marconi *et al*, 2021).

**Movie S1.**

Time-lapse video of the root apex of a wild type seedling of the cell cycle marker PlaCCI line.

**Movie S2.**

Time-lapse video of the root apex of a *plt1-4,plt2-2* mutant seedling expressing the PlaCCI cell cycle markers.

**Movie S3.**

Time-lapse video of the root apex of a *krp5-1* mutant seedling expressing the PlaCCI cell cycle markers.

**Movie S4.**

Time-lapse video of the root apex of a *rRBr* seedling (a RNAi line directed to RBR1 in the root apical meristem) expressing the PlaCCI cell cycle markers.

**References**

Brady SM, Orlando DA, Lee JY, Wang JY, Koch J, Dinneny JR, Mace D, Ohler U & Benfey PN (2007) A high-resolution root spatiotemporal map reveals dominant expression patterns. *Science* 318: 801–806

Desvoyes, B., Echevarria, C., & Gutierrez, C. (2021) A perspective of cell proliferation kinetics in the root apical meristem. *J Exp Bot* 72: 6708-6715

Marconi M, Gallemi M, Benková E & Wabnik K (2021) A coupled mechano-biochemical framework for root meristem morphogenesis. *bioRxiv*: 2021.01.27.428294

Pacheco-Escobedo MA, Ivanov VB, Ransom.Rodriguez I, Arriaga-Mejía G, Avila H, Baklanov IA, Pimentel A, Corkidi G, Doerner P, Dubrovsky JG, *et al* (2016) Longitudinal zonation pattern in Arabidopsis root tip defined by multiple structural change algorithm. *Ann Bot* 118: 763–776

Salvi E, Rutten JP, Di Mambro R, Polverari L, Licursi V, Negri R, Dello Ioio R, Sabatini S & Ten Tusscher K (2020) A Self-Organized PLT/Auxin/ARR-B Network Controls the Dynamics of Root Zonation Development in Arabidopsis thaliana. *Dev Cell* 53: 431-443.e23

Scheres B (2007) Stem-cell niches: nursery rhymes across kingdoms. *Nature reviews* 8: 345–54

Shimotohno A & Scheres B (2019) Topology of regulatory networks that guide plant meristem activity: similarities and differences. *Curr Opin Plant Biol* 51: 74–80

Svolacchia N, Salvi E & Sabatini S (2020) Arabidopsis primary root growth: let it grow, can't hold it back anymore! *Curr Opin Plant Biol* 57: 133–141
