## Supplementary Table S2 for "Stem cell regulators control a G1 duration gradient in the plant root meristem"

Supplementary Table 1

| **Parameter** | **Description** | **Value** | **Units** |
| --- | --- | --- | --- |
| *PR_max_* | maximum level of auxin-induced *driver* expression | 20 | nM/h |
| *PR_k_* | half-max coefficient of auxin-induced *driver* expression | 2 | nM/μm^2^ |
| *PR_perm_* | *driver* cell permeability coefficient | 1 | μm/h |
| *d_PR_* | *driver* degradation rate | 0.01 | h^-1^ |
| *IN_max_* | maximum level of *driver*-induced *regulator* expression | 20 | nM/h |
| *IN_k_* | half-max coefficient of *driver*-induced *regulator* expression | 5 | nM/μm^2^ |
| *d_IN_* | *regulator* degradation rate | 0.01 | μm/h |
| *DIV_max_* | maximum cell cycle time | 50 | h |
| *DIV_IN_* | exponential rate of cell cycle regulation by the *regulator* | 0.03 | h^-1^ |

For a global description of the default parameters please refer to the original publication <https://www.biorxiv.org/content/10.1101/2021.01.27.428294v2>
